# ACOD1 regulates microglial arginine metabolism and inflammatory responses

**DOI:** 10.1101/2025.06.01.657243

**Authors:** Eleftheria Karadima, Canelif Yilmaz, Anupam Sinha, Georgia Fodelianaki, Sofia Dimothyra, Nikolaos Nirakis, Sofia Traikov, Nicola Zamboni, Ben Wielockx, Panayotis Verginis, Mirko Peitzsch, Triantafyllos Chavakis, Vasileia Ismini Alexaki

## Abstract

Itaconate is produced by inflammatory macrophages and promotes a negative feedback on inflammation. It is synthesized by aconitate decarboxylase 1 (ACOD1) from cis-aconitate, a metabolite of the tricarboxylic acid cycle. Here, we studied the role of ACOD1 in the inflammatory response of microglia, the resident macrophage-like cells of the brain. Similar to macrophages, ACOD1 deficient microglia displayed a stronger inflammatory response to lipopolysaccharide (LPS) compared to their wild type counterparts. The proinflammatory effects of ACOD1 deficiency were associated with reprogramed arginine metabolism entailing enhanced argininosuccinate synthesis at the expense of polyamine synthesis, in a manner that was dependent on ACLY. These findings provide new insights in the immunometabolic role of ACOD1 in inflammatory microglia.

## Introduction

Macrophage immune responses are orchestrated by cell metabolic reprograming (1). Itaconate, a byproduct of the tricarboxylic acid cycle (TCA), is produced in inflammatory macrophages and negatively feedbacks on inflammation (2). It is synthesized through decarboxylation of cis-aconitate by aconitate decarboxylase 1 (ACOD1, encoded by Immune responsive gene 1 (*Irg1* or *Acod1*)), the expression of which is induced in macrophages by inflammatory stimuli, such as lipopolysaccharide (LPS) (3). Itaconate reprograms the TCA cycle by inhibiting succinate dehydrogenase (SDH), leading to accumulation of succinate (4, 5). Several molecular mechanisms mediate the anti-inflammatory effects of itaconate including 1) alkylation of KEAP1 and downstream activation of Nuclear factor erythroid 2-related factor 2 (NRF2), 2) alkylation-mediated inhibition of stimulator of interferon genes (STING), 3) Activating Transcription Factor 3 (ATF3)-mediated induction of IκBζ and 4) inhibition of TET-family DNA dioxygenases and thereby downregulation of NF-κB and STAT target genes (6–9). Consequently, itaconate reduces the expression of pro-inflammatory cytokines, such as IL-1b, and IL-6, and downregulates production of reactive oxygen species (ROS) (5). The anti-inflammatory function of the ACOD1-itaconate axis was shown in infection, sepsis, myocardial disease, atherosclerosis, autoimmune disease and gout (7, 8, 10–15).

Microglia are the resident macrophage-like cells of the central nervous system (CNS) (16). They assist in proper synaptic remodeling (17) and maintain brain homeostasis via removal of damaged or dead cells and debris (18). In neurodegenerative diseases, microglia lose their homeostatic function and acquire an inflammatory phenotype (19). Sustained low-grade microglia inflammation is a common feature of many neurological diseases, such as Alzheimer’s disease and multiple sclerosis (MS) (20). However, the mechanisms and particularly the cell metabolic circuits regulating microglia-mediated neuroinflammation are still inadequately understood. Here, we investigated the role of ACOD1 in microglial inflammatory responses. We show that LPS treatment induces ACOD1 expression and itaconate production in microglia. ACOD1 deficiency increased ATP citrate lyase (ACLY) expression and acetyl-CoA levels and reprogramed arginine metabolism towards argininosuccinate synthesis at the expense of polyamine metabolism. We suggest that ACOD1 functions as a regulatory metabolic checkpoint for cell metabolism fine-tuning inflammatory responses in microglia.

## Materials and Methods

### Mice and in vivo experiments

*Acod1^-/-^* mice were purchased from The Jackson Laboratory (JAX #029340). Wild type (wt) C57BL/6J mice were obtained from Charles River Laboratories. Eight to twelve-week old male mice were intraperitoneally (i.p.) injected with 3 mg/kg LPS (LPS-EB Ultrapure; InVivoGen, tlrl-3pelps) or PBS and after 4, 16 or 24 hours they were sacrificed by cervical dislocation. In other experiments, wt C57BL/6J mice were i.p. injected with 50 mg/kg argininosuccinate, or PBS and after 3 h 1 mg/kg LPS was i.p. injected. Four h later mice were killed by cervical dislocation. All animal experiments were in compliance with the local ethical guidelines and approved by Landesdirektion Sachsen, Germany.

### Microglia isolation, culture and treatments

Primary microglia were isolated as previously described (21, 22). Briefly, mouse brains of 8-9 week-old wt and *Acod1*^-/-^ mice were digested with an enzymatic solution containing 0.5 mM EDTA, 5 mM L-cysteine (Sigma-Aldrich), 0.1 mg/ml papain (Sigma-Aldrich,) and 2.4 mg/ml dispase II (Sigma Aldrich) diluted in DMEM (Thermo Fisher Scientific). The enzymatic reaction was stopped by adding 20% FBS in PBS. After centrifugation at 1,200 rpm for 7 min at 4 °C, cells were re-suspended in 0.5 mg/ml DNase I (Thermo Fisher Scientific) in PBS and incubated for 5 min in room temperature (RT). Cells were gently dissociated and passed through a 100 μm cell strainer. Isolated cells were cultured in DMEM/F12 (Thermo Scientific) with Glutamax, 10% FBS, 1% penicillin/streptomycin (P/S) and 10 ng/ml granulocyte and macrophage colony stimulating factor (GM-CSF) (Peprotech) in poly-L-Lysine-coated flasks and maintained in culture at 37 °C and 5% CO2. Cells were treated with LPS (100 ng/ml, tlrl-peklps, InvivoGen), IFNg (20 ng/ml, Thermo Fisher Scientific), 4-octyl itaconate (4-OI, 125 μM, Sigma-Aldrich), BMS303141 (20 μM, Sigma-Aldrich), spermidine (10 μM, Sigma-Aldrich) or respective vehicle controls.

### Isolation of glial populations and neurons

Mouse brains were dissected and the Neural Tissue Dissociation Kit (Miltenyi Biotec) was used for obtaining single-cell suspensions according to manufacturer’s instructions. Sequential isolation of microglia, oligodendrocytes and astrocytes from the same samples was achieved by positive selection after serial incubation with anti-CD11b Microglia Microbeads (Miltenyi Biotec), anti-O4 Microbeads (Miltenyi Biotec) and anti-astrocyte cell surface antigen-2 (ACSA-2) Microbeads kit (Miltenyi Biotec) with sequential passage through LS columns. Neurons were isolated by negative selection for CD11b, O4 and ACSA-2.

### Bone marrow myeloid cell ex vivo treatment with LPS

Bone marrow of wt male mice was isolated from femur and tibia by flushing the bones with 10 ml of 0.5% BSA in PBS supplemented with 2 mM EDTA. The cell suspension was centrifuged at 500xg for 5 min at RT, followed by a 3-min incubation with red blood cell (RBC) lysis buffer (eBioscience). Following RBC lysis, the cell suspension was passed through a 40 μm cell strainer and then centrifuged at 500xg for 5 min at RT and the cell pellet was resuspended in the appropriate volume of RPMI 1640 medium, supplemented with 10% FBS and 1% P/S. The cells were incubated at 37 °C for 1 h to attach and were then treated for 4 h with 100 ng/ml LPS or carrier (PBS). Afterwards, they were scraped, centrifuged and stained with anti-CD45-PerCPCy5.5 (1:100, 103132, Biolegend), anti- CD11b-FITC (1:100, 101206, Biolegend) or anti-Ly6G-APC (1:100, 560599, BD Biosciences) in 5% FBS in PBS for 20 min at 4 °C in the dark. CD45^high^CD11b^+^Ly6G^-^ myeloid cells and neutrophils (CD45^high^CD11b^+^Ly6G^+^ cells) were sorted using a BD FACSAria II (BD Biosciences) and the FACSDiva software (BD Biosciences). Sorted cells were collected in 10% FBS in PBS and centrifuged at 1,400 rpm for 10 min at 4 °C and the cell pellet was lysed in RA1 buffer with beta-mercaptoethanol for further RNA isolation.

### Sorting of spleen T cells and myeloid cells

Spleens were isolated from PBS or LPS treated mice and smashed in 5 ml of 1% FBS in PBS supplemented with 2 mM EDTA through a pre-wet 100 μm cell strainer. Cells were centrifuged at 1,500 rpm for 5 min at 4 °C, incubated with RBC lysis buffer for 3 min at RT and then washed with 10 ml of 1% FBS in 1x PBS supplemented with 2 mM EDTA. The cells were stained with anti-CD45-PerCPCy5.5 (1:100, 103132, Biolegend), anti-CD11b- FITC (1:100, 101206, Biolegend), anti-Ly6G-APC (1:100, 560599, BD Biosciences), anti-CD3-PE (1:100, 130-102-600, Milteniy Biotec), anti-CD4-APC (1:100, 130-116-546, Milteniy Biotec), or anti-CD8a-PerCP (1:100, 130-122-953, Milteniy Biotec). CD45^high^CD11b^+^Ly6G^-^ myeloid cells, T helper cells (CD3^+^CD4^+^) and cytotoxic T cells (CD3^+^CD8^+^ cells) were sorted using a BD FACSAria II (BD Biosciences) and the FACSDiva software (BD Biosciences).

### FACS analysis and sorting of brain microglia and monocyte/macrophages

Brains were smashed in isolation buffer (0.5% BSA PBS) on ice and cell suspensions were centrifuged at 300 x g for 10 min at 4 °C. Myelin was removed with Myelin Removal Beads II (Miltenyl Biotec) and LS columns (Miltenyl Biotec) per manufacturer’s instructions. The cells were incubated with anti-CD45-PerCPCy5.5 (1:100, 103132, Biolegend), anti-CD11b-FITC (1:100, 101206, Biolegend) or anti-Ly6G-APC (1:100, 560599, BD Biosciences) in 5% FBS PBS for 30 min at 4 °C in the dark. FACS analysis was performed using a BD FACSCanto II flow cytometer (BD Biosciences) and the FACSDiva 6.1.3 software (BD Biosciences). Microglia were identified as CD45^interm^CD11b^+^Ly6G^-^ and monocytes/macrophages as CD45^high^CD11b^+^Ly6G^-^. Microglia were sorted with a BD FACSAria II (BD Biosciences) and the FACSDiva software (BD Biosciences). Sorted cells were collected in 10% FBS in PBS, centrifuged at 1,400 rpm for 10 min at 4 °C and the cell pellet was stored at -80 °C until further analysis.

### BV2 cell culture and treatments

BV2 cells were obtained from Interlab Cell Line Collection (ICLC, Genova, Italy) and maintained in RPMI-1640 medium supplemented with 10% FBS and 1% P/S at 37 °C and 5% CO2. BV2 cells were treated for 4 h with following TLR ligands: PAM3CSK4 (TLR1/TLR2 ligand, 1 μg/ml), heat-killed preparation of Listeria monocytogenes (HKLM, TLR2 ligand, 10^8^ cells/ml), polyinosinic-polycytidylic acid (poly(I:C), TLR3 ligand, 1 μg/ml), flagellin from Salmonella Typhimurium (FLA-ST, TLR5 ligand, 1 μg/ml), FSL-1 (TLR2/6 ligand, 100 ng/ml), imiquimod (TLR7 ligand, 1 μg/ml), ssRNA40/Lyovec (TLR8 ligand, 1 μg/ml) and Class B CpG oligonucleotide (ODN2006, TLR9 ligand, 5 μM), all from the Human TLR1-9 Agonist kit (InvivoGen, tlrl-kit1hw). Also, BV2 cells were treated for 4 h with M-CSF, GM-CSF, interleukin 1b (IL-1b), IL-6, tumor necrosis factor (TNF), IL-4, IL- 10, TGF-b or IFNg (all at 20 ng/ml from Peprotech).

### siRNA transfections

Primary microglia cells were transfected with small interfering RNAs (siRNAs) and Lipofectamine™ RNAiMAX Transfection Reagent (Invitrogen) using the forward transfection protocol according to the manufacturer’s protocol. Cells were incubated with 30 nM siRNAs for 24 h (si*Ass1* and siControl) or 48 h (si*Odc* and siControl). All siRNAs were purchased from Dharmacon-Horizon Discovery.

### RNA-seq

Bulk RNA-seq was performed and analyzed as previously described (23, 24). For transcriptome mapping, strand-specific paired-end sequencing libraries from total RNA were constructed using TruSeq stranded Total RNA kit (Illumina Inc). Sequencing was performed on an Illumina HiSeq3000 (1x75 basepairs). Low quality nucleotides were removed with the Illumina fastq filter and reads were further subjected to adaptor trimming using cutadapt (25). Alignment of the reads to the mouse genome was done using STAR Aligner (26) using the parameters: “–runMode alignReads –outSAMstrandField intronMotif –outSAMtype BAM SortedByCoordinate --readFilesCommand zcat”. Mouse Genome version GRCm38 (release M12 GENCODE) was used for the alignment. The parameters: ‘htseq-count -f bam -s reverse -m union -a 20’, HTSeq-0.6.1p1 (27) were used to count the reads that map to the genes in the aligned sample files. The GTF file (gencode.vM12.annotation.gtf) used for read quantification was downloaded from Gencode (https://www.gencodegenes.org/mouse/release_M12.html). Gene centric differential expression analysis was performed using DESeq2_1.8.1 (28). The raw read counts for the genes across the samples were normalized using ‘rlog’ command of DESeq2 and subsequently these values were used to render a PCA plot using ggplot2_1.0.1 (29).

Pathway and functional analyses were performed using GSEA (29) and EGSEA (30). GSEA is a stand-alone software with a graphical user interface (GUI). To run GSEA, a ranked list of all the genes from DESeq2 based calculations was created using the -log10 of the p-value. This ranked list was then queried against Molecular Signatures Database (MsigDB), KEGG, GO, Reactome and Hallmark based repositories. EGSEA is an R/Bioconductor based command-line package. For pathway analysis with EGSEA lists of DEG with padj < 0.05 or p < 0.05 were used based on the KEGG database repository. For constructing heatmaps, the “rlog-normalized” expression values of the significantly expressed genes (padj < 0.05) was scaled using z-transformation. The resulting matrices were visually rendered using MORPHEUS (24, 31).

### RNA isolation and quantitative RT-PCR

Total RNA was extracted from cells or tissues using the Nucleospin RNA isolation kit (Macherey-Nagel), according to manufacturer’s instructions. cDNA was synthesized using the iScript cDNA synthesis kit (Bio-Rad). qPCR was performed using the SsoFast Eva Green Supermix (Bio-Rad), a CFX384 real-time System C1000 Thermal Cycler (Bio- Rad), and the Bio-Rad CFX Manager 3.1 software. The relative amount of mRNA was calculated with the ΔΔCt method, using *18s* as a housekeeping gene. The primer sequences are listed in Table 1.

### Non-targeted metabolomics

Cells were washed with 75 mM ammonium carbonate at pH 7.4 and cell pellets were collected and frozen in liquid nitrogen. Intracellular metabolites were extracted twice with 70% ethanol at 75°C for 3 min, dried in a speedvac and resuspended in H2O. Extracts were analyzed by flow injection – time of flight mass spectrometry on an Agilent 6550 QTOF instrument, as described previously (32). Ion annotation was based on matching their measured masses to that of the compounds listed in the KEGG mmu database.

### Targeted metabolomics

TCA metabolites were measured as previously described (33). Briefly, metabolites were extracted from samples with methanol, dried, resuspended in mobile phase and cleared with a 0.2 µm centrifugal filter. To improve separation, the elution gradient was changed as follows: 99% A (0.2% formic acid in water), 1% B (0.2% formic acid in acetonitrile) for 2.00 min, 100% B at 2.50 to 2.65 min, 1% B at 3.40 min and equilibration with 1% B until 5.00 min. Multiple reaction monitoring with negative electrospray ionization was used for quantification. Itaconate was measured using multi-reaction monitoring (MRM)-derived ion transition of 128.9→85.1. For quantification of itaconate ratios of analyte peak areas to respective peak areas of the stable isotope labeled internal standard (itaconic acid- ^13^C5; Bio-Connect B.V., The Netherlands; MRM transition 133.9→89.1) obtained in samples were compared to those of calibrators.

Arginine metabolites were measured as previously described (34). Sample preparation was performed by addition of 10 µl internal standard working solution followed by 200 µL H2O:acetonitrile 50:50 (v/v) extraction buffer and subsequent grinding for 30 sec. After homogenization, samples were vortex-mixed for one minute and centrifuged at 3,000g for 10 min at 4°C. Clear supernatants were transferred directly onto a 96-well- polytetrafluoroethylene (PTFE)-filterplate (Merck-Millipore) and filtered by assistance of positive pressure. Subsequently, filtered extracts were dried in a vacuum-assisted centrifuge, thereafter reconstituted in 200 µl initial mobile phase and analyzed by LC-MS/MS. LC-MS/MS measurements were performed on a QTRAP® 6500+ triple quadrupole mass spectrometer from Sciex coupled to a Waters Acquity ultra-performance liquid chromatography system. Chromatographic separation was achieved by using a XBridge BEH Amide XP Column (2.1 x 100 mm, 2.5 µm; Waters) at 40 °C using a gradient of mobile phases A (20mM ammonium formate / 5% methanol at pH 3) and B (ACN/methanol/mobile phase A, 90%/5%/5%). Five µL of reconstituted calibrators, QC samples and test samples, kept at 4°C in the autosampler, were injected into the LC-MS/MS system at a flow rate of 0.4 mL/min with 15% mobile 29 phase A. At 0.37 min, mobile phase A started to linearly increase up to 30% at 4.1 min and further to 50% at 5 min. At 5.8 min, mobile Phase A increased up to 85% and after a hold until 6.8 min, the gradient returned back to initial conditions at 7.8 min, followed by another 1.7 min for column re-equilibration

### Acetyl-CoA measurement

Acetyl-CoA was measured as previously described (23). Cultured cells were washed with cold PBS, scraped on ice in cold PBS, centrifuged for 2 min at 1,500 rpm at 4 °C, washed once with 0.1 M ammonium bicarbonate, and centrifuged again for 2 min at 1,500 rpm at 4 °C. Cell pellets and tissues were snap-frozen and stored at -80 °C until further analysis.

Frozen samples were dissolved in 150 μl of 30% methanol in acetonitrile containing 100 nM AMP – isotope-labeled (Adenosine-¹³C₁₀,¹⁵N₅-5′-monophosphate) as an internal standard. LC–MS/MS analysis was performed using a high-performance liquid chromatography (HPLC) system (Agilent 1200) coupled online to a G2-S QTof mass spectrometer (Waters). For normal-phase chromatography, Bridge Amide 3.5 μm (2.1 × 100 mm) columns (Waters) were used. The mobile phase consisted of eluent A (95% acetonitrile, 0.1 mM ammonium acetate, and 0.01% NH₄OH) and eluent B (40% acetonitrile, 0.1 mM ammonium acetate, and 0.01% NH₄OH), applied with the following gradient program: 0% to 100% eluent B within 18 min, 100% eluent B from 18 to 21 min and 0% eluent B from 21 to 26 min. The flow rate was set to 0.3 ml/min. The spray voltage was set to 3.0 kV, and the source temperature was maintained at 120 °C. Nitrogen was used as both the cone gas (50 l/h) and desolvation gas (800 l/h), while argon was used as the collision gas. The MSE mode was applied in negative ionization polarity. Mass chromatograms and spectral data were acquired and processed using MassLynx software (Waters).

### Western blotting

Protein extracts were prepared in ice-cold RIPA lysis buffer system supplemented with protease and phosphatase inhibitors (SCBT) or lysis buffer supplemented with PhosSTOP (Roche) and cOmplete^™^, Mini, EDTA-free Protease Inhibitor Cocktail (Roche) Protein concentration was determined with the BCA assay (Thermo Fisher Scientific). Protein lysates were mixed with reducing Laemmli SDS sample buffer (Thermo Fisher Scientific), denatured at 95 °C for 5 min and loaded on a polyacrylamide gel and separated with SDS-PAGE. Afterward, proteins were transferred onto nitrocellulose membranes and blocking was performed with 5% BSA TBS-T buffer for 1 h at RT followed by overnight incubation with the primary antibody. Primary antibodies used were following: anti-ACOD1 (Abcam, ab222411), anti-IL-1b (Cell Signaling Technology, #12507), anti-phosphoACLY (Ser455) (Cell Signaling Technology, #4331T), anti-Vinculin (Cell Signaling Technology, #4650), anti-Tubulin (Sigma-Aldrich, T5186) and anti-b-actin (Cell Signaling Technology, #4970) all diluted at 1:1,000 in 5% BSA TBS-T. Next, goat anti-rabbit IgG horseradish peroxidase-conjugated antibody (1:3,000, R&D Systems, HAF008) was added to the membranes and incubated for 2 h at RT. Finally, membranes were washed with TBS-T and developed using SuperSignal West Pico Chemiluminescent Substrate (Life Technologies) or SuperSignal West Fempto Chemiluminescent Substrate (Life Technologies) and a LAS-3000 luminescent image analyzer (Fujifilm). The intensity of the bands was quantified using the FIJI software.

### ELISA

For the quantification of IL-6 in cell culture supernatants, mouse IL-6 DuoSet ELISA (#DY406-ML, R&D Systems) was used according to manufacturer’s instructions.

### Statistical analyses

The statistical analysis and data plotting were done with the GraphPad Prism 10 software. All values are expressed as mean ± SEM. Data were analyzed with Student’s t-test if normally distributed, Mann Whitney U-test if non-normally distributed; One-way analysis of variance (ANOVA) with post hoc Tukey’s test for multiple comparisons. p < 0.05 or adjp < 0.05 were set as significance levels.

## Results

### Inflammation induces itaconate production in microglia

First, we validated that inflammation induces *Acod1* expression in microglia. Wt C57BL/6J mice were treated i.p. for 4 h with LPS and whole brain microglia were FACS sorted as CD45^interm^CD11b^+^Ly6G^-^ cells, distinguished from monocytes/macrophages (CD45^high^CD11b^+^Ly6G^-^) and neutrophils (CD45^high^CD11b^+^Ly6G^+^). Transcriptional changes in microglia from LPS- and PBS-treated mice were assessed by bulk RNA-seq (Figure 1A). In total, 3,407 genes were upregulated and 3,367 genes were downregulated in microglia of LPS-treated mice (Figure 1A). Microglia of LPS-treated mice exhibited strong enrichment of inflammatory response-related gene sets, as shown by gene set enrichment analysis (GSEA) (Figure 1B). LPS treatment strongly induced *Acod1* expression, while it reduced the expression of several genes encoding for proteins associated with the TCA cycle, such as isocitrate dehydrogenase 1 (*Idh1*), *Idh3g*, succinate-CoA ligase GDP/ADP-forming subunit alpha (*Suclg1*), malate dehydrogenase 1 (*Mdh1*) and aconitase 1 (*Aco1*) (Figure 1A,C). In accordance, previous studies reported that inflammation induced TCA cycle fragmentation by reducing the expression of SDH and IDH (4, 5, 24, 35).

**Figure 1.**
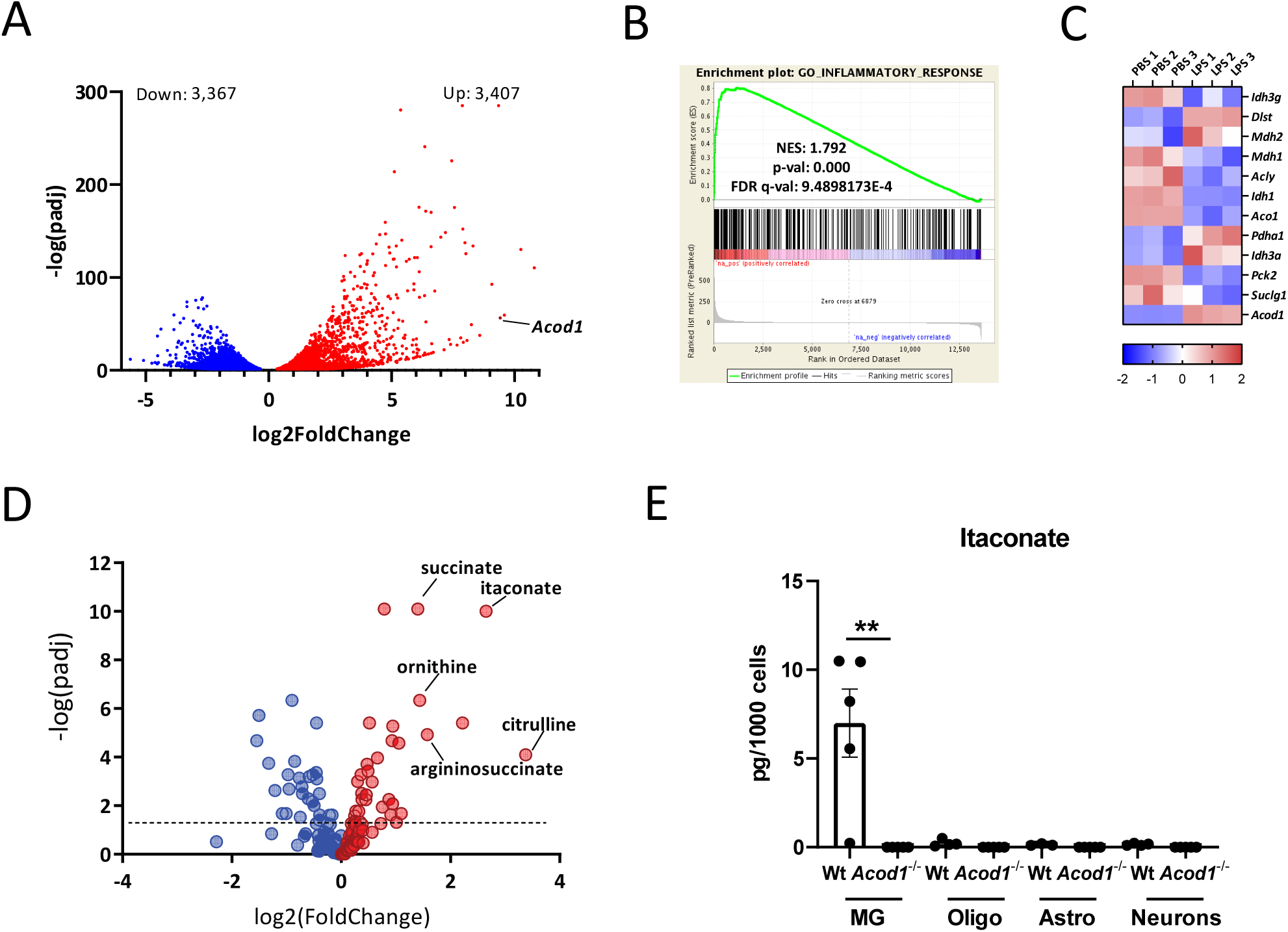
Inflammation induces itaconate production in microglia. (A-C) Bulk RNA- seq in sorted microglia (CD45^interm^CD11b^+^Ly6G^-^) from wt mice i.p. treated for 4 h with PBS or LPS (3 mg/kg)(n=3 mice per group). **(A)** Volcano plot showing differentially expressed genes. **(B)** Gene set enrichment analysis (GSEA) for inflammatory response- related genes. NES, normalized enrichment score; FDR, false discovery rate. **(C)** Heatmap of differentially expressed TCA cycle-related genes (padj < 0.05) from LPS- treated wt mice. **(D)** Volcano plot showing regulated metabolites in primary microglia treated or not with LPS (100 ng/ml) for 24 h (n=5 biological samples per group). **(E)** Itaconate amounts in isolated microglia (MG), oligodendrocytes (Oligo), astrocytes (Astro) and neurons from brains of wt and *Acod1*^-/-^ mice treated for 24 h with LPS (n=5 mice per group). **p < 0.01.

Similarly, *Acod1* was amongst the most upregulated genes in mouse primary microglia treated for 4 h with LPS, assessed by bulk RNA-seq analysis (Figure S1A). LPS-treated microglia showed enrichment of inflammatory response-related gene sets (Figure S1B), while expression of several TCA cycle-related genes, such as succinate dehydrogenase b (*Sdhb*), *Sdhd*, *Idh3g*, *Idh1*, *Idh2*, *Idh3b* and *Suclg2* was downregulated (Figure S1C). Upregulation of *Acod1* gene and protein expression was verified in primary microglia treated with LPS+IFNg for 4 and 24 h, respectively (Figure S1D,E).

Accordingly, itaconate and succinate were the most upregulated metabolites in primary microglia treated for 24 h with LPS, as shown by non-targeted metabolomics (Figure 1D).

Succinate accumulation was in accordance with the inhibitory effect of itaconate on SDH and reduced expression of *Sdhb*, *Sdhd*, *Suclg1* and *Suclg2* in inflammatory microglia (Figure 1C,S1C) (5, 24). Moreover, LPS treatment led to increased intracellular amounts of arginine metabolites, including ornithine, citrulline and argininosuccinate (Figure 1D). Increased citrulline and argininosuccinate levels indicate activation of the arginine biosynthesis pathway sustaining nitric oxide production in inflammatory macrophages (36, 37). Increased ornithine amounts are due to upregulation of arginase 1 and 2 (ARG1, ARG2) and facilitate polyamine and proline synthesis that may be involved in resolution of inflammation and tissue recovery in the later stages of the inflammatory response (36).

Next, we asked which cell types produce itaconate in the brain upon inflammation. Monocytes and neutrophils were previously reported to express *Acod1* upon inflammation or infection (38–40). We verified that LPS induced *Acod1* expression in bone marrow CD45^high^CD11b^+^Ly6G^-^ myeloid cells and neutrophils (Figure S2A). Moreover, spleen CD4^+^ and CD8^+^ T cells, in contrast to spleen CD45^high^CD11b^+^Ly6G^-^ myeloid cells, did not express *Acod1* in response to LPS (Figure S2B). Spleen CD45^high^CD11b^+^Ly6G^-^ myeloid cells and whole brain microglia of LPS-treated mice contained similar amounts of itaconate (Figure S2C). Of note, monocytes/macrophages were very few compared to microglia in the brains of LPS-treated perfused mice (Figure S2D), suggesting that microglia are a more important source of itaconate in the brain of LPS-treated mice compared to monocytes/macrophages.

Moreover, we examined whether brain cell types other than microglia may produce itaconate. To this end, microglia and brain resident macrophages (CD11b^+^), oligodendrocytes (O4^+^), astrocytes (ACSA-2^+^) and neurons (negative for CD11b, O4 and ACSA-2) were sorted from wt and *Acod1^-/-^* mice treated for 24 h i.p. with LPS, and analyzed by LC-MS/MS (Figure S2E). Itaconate was detected in high amounts in CD11b^+^ cells (microglia/brain macrophages) but not in the other cell populations (oligodendrocytes, astrocytes, neurons), and its production was completely blunted in ACOD1 deficient mice (Figure 1E). Similarly, LPS-induced *Acod1* expression was fully suppressed in microglia sorted from *Acod1^-/-^* mice (Figure S2F) and LPS+IFNg-induced itaconate production was completely blunted in *Acod1^-/-^* primary microglia (Figure S2G).

Finally, we asked which inflammatory stimuli induce *Acod1* expression in microglia. To this end, BV2 cells were treated for 4 h with different Toll like receptor (TLR) ligands, such as PAM3CSK4 (TLR1/TLR2 ligand), HKLM (TLR2 ligand), poly(I:C) (TLR3 ligand), FLA-ST (TLR5 ligand), FSL1 (TLR2/6 ligand), imiquimod (TLR7 ligand), ssRNA40/Lyovec (TLR8 ligand) and ODN2006 (TLR9 ligand), different cytokines, like M-CSF, GM-CSF, IL- 1b, IL-6, TNF, IL-4, TGF-b, IL-10, IFNg, and LPS or LPS+IFNg, and *Acod1* expression was examined by qPCR. Out of the tested substances, LPS most strongly induced *Acod1* expression, but PAM3CSK4, HKLM, polyI:C, FSL1, ODN2006, IL-1b, TNF and IFNg also upregulated *Acod1* expression (Figure S3).

### ACOD1 knockout enhances the inflammatory response of microglia

Next, we examined the role of ACOD1 in microglia-mediated inflammation. To this end, *Acod1^-/-^* and littermate wt mice were treated for 16 h with LPS, and whole brain microglia were sorted as CD45^interm^CD11b^+^Ly6G^-^ cells and analyzed by bulk RNA-seq. In total 309 genes were upregulated and 261 genes were downregulated in microglia of *Acod1^-/-^* compared to wt mice (Figure 2A). Upregulated genes included mediators of inflammation, such as interleukin 1 receptor type 2 (*Il1r2)*, interleukin 6 receptor subunit alpha (*Il-6ra)*, secreted phosphoprotein 1 (*Spp1)*, *Arg1*, Triggering Receptor Expressed On Myeloid Cells 1 (*Trem1)*, Sphingosine Kinase 1(*Sphk1)*, Toll Like Receptor 5 (*Tlr5)*, Complement C5a Receptor 2 (*C5ar2)*, Cytotoxic And Regulatory T Cell Molecule (*Crtam)*, Interferon Regulatory Factor 4 (*Irf4)* and Cadherin 11 *(Cdh11)* (Figure 2A). Accordingly, GSEA analysis showed significant positive enrichment of gene sets related to the innate immune system in microglia of *Acod1^-/-^* compared to wt mice (Figure 2B). ACOD1 deficiency increased IL-1b and IL-6 mRNA and protein expression in inflammatory microglia (Figure 2C-E), standing in accordance with its reported effects in LPS-treated macrophages (5). Moreover, in accordance with these reports in macrophages, ACOD1 deficiency did not increase LPS-induced *Tnf* expression in microglia (not shown) (5). Moreover, the effects of ACOD1 deficiency on *Il-1b* and *Il-6* expression were completely blunted by 4-OI, a membrane-permeable derivative of itaconate (Figure 2E). In accordance with the inhibitory effect of itaconate on SDH (5), ACOD1 deficiency reduced intracellular succinate amounts and the succinate/fumarate ratio in inflammatory microglia (Figure S4A,B).

**Figure 2.**
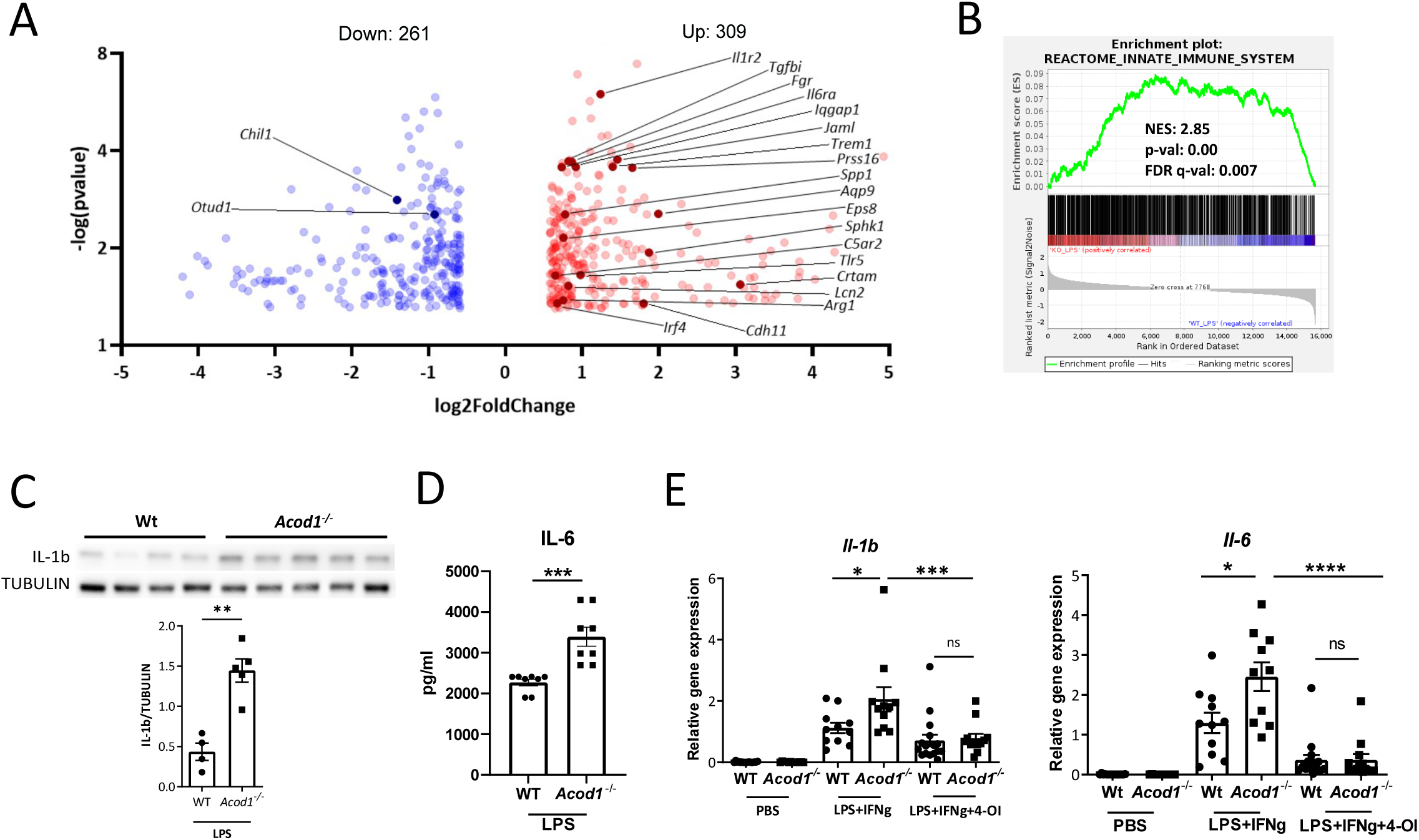
ACOD1 knockout enhances the inflammatory response of microglia. (A,B) Bulk RNA-seq in sorted microglia (CD45^interm^CD11b^+^Ly6G^-^) from *Acod1*^-/-^ and wt mice treated for 16 h with LPS (3 mg/kg) (n=3 mice per group). **(A)** Volcano plot showing differentially expressed genes. **(B)** GSEA for innate immune system-related genes. **(C)** Western blot analysis for IL-1b after treatment of *Acod1*^-/-^ and wt microglia for 24 h with LPS+IFNg and band intensity quantification using Tubulin as a loading control (n=4-5). **(D)** IL-6 amounts in supernatants of *Acod1^-/-^* and wt microglia treated for 4 h with LPS (n=8). **(E)** *Il-1b* and *Il-6* expression in *Acod1*^-/-^ and wt microglia treated for 24 h with LPS+IFNg, 4-OI (125 μM) or respective carriers (n=10-16). *p < 0.05, **p < 0.01, ***p < 0.001, ****p < 0.0001, ns: non-significant.

### ACOD1 deficiency reprograms arginine metabolism

Arginine metabolism is a key metabolic hub in the regulation of macrophage immune responses (36). Proinflammatory macrophages metabolize arginine to nitric oxide and citrulline, and in turn, citrulline can be used to regenerate arginine in order to sustain nitric oxide production (36, 37). This requires the conversion of citrulline to argininosuccinate by argininosuccinate synthase 1 (ASS1) and the break-down of argininosuccinate to arginine and fumarate by the argininosuccinate lyase (ASL) (36). The citrulline-nitric oxide cycle is activated in inflammatory macrophages and ASS1-mediated intracellular citrulline depletion is required for the proinflammatory response of macrophages (37, 41). We showed that citrulline and argininosuccinate accumulate in LPS-stimulated microglia (Figure 1D). Interestingly, ACOD1 deficiency enhanced argininosuccinate amounts in LPS-stimulated microglia (Figure 3A). Upregulation of argininosuccinate production in ACOD1 deficient microglia could be due to release of SDH inhibition (Figure S4), permitting metabolism of fumarate via malate to aspartate, the latter, together with citrulline, being the substrate of ASS1 (42). Accordingly, microglia sorted from brains of LPS-treated *Acod1^-/-^* mice displayed increased *Ass1* expression compared to microglia of wt mice (Figure 3B) and siRNA silencing of *Ass1* reduced *Il-1b* expression in *Acod1*^-/-^ microglia (Figure 3C). In order to validate the proinflammatory role of argininosuccinate *in vivo*, we treated wt mice i.p. with argininosuccinate prior to LPS treatment and brain microglia were sorted 4 h after the LPS injection. Argininosuccinate treatment of LPS- treated mice further increased *Il-1b* expression and enhanced the LPS-mediated suppression of the homeostatic genes *Sall1*, *Tmem119* and *Cx3cr1*, and phagocytic genes *Trem2* and Mer tyrosine kinase (*Mertk*) (Figure 3D)(43, 44). These data suggest that ACOD1 deficiency enhances argininosuccinate production, which promotes IL-1b- mediated inflammation and reduces homeostatic microglia features.

**Figure 3.**
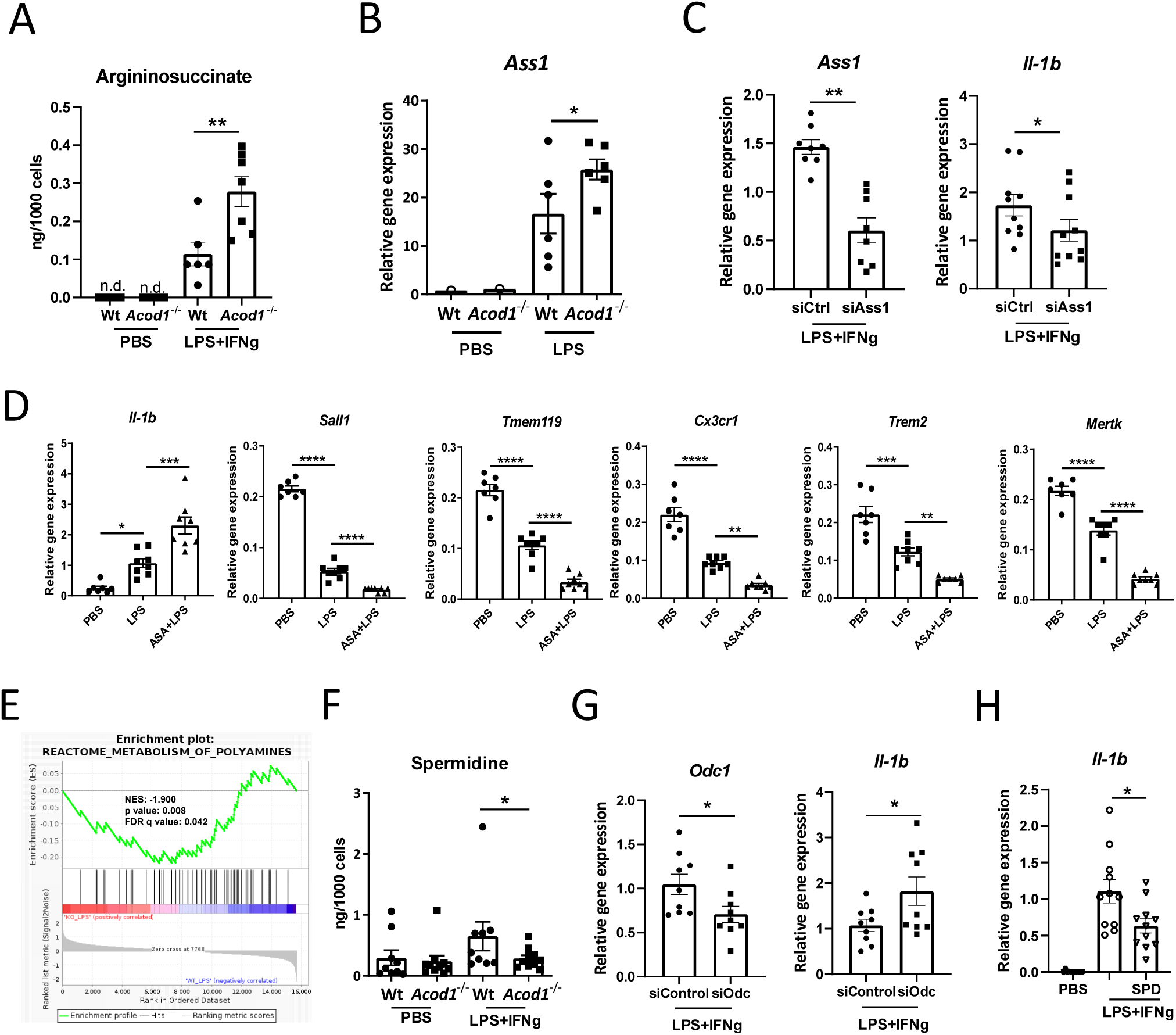
ACOD1 deficiency reprograms arginine metabolism. **(A)** Argininosuccinate amounts in primary *Acod1*^-/-^ and wt microglia treated for 24 h with LPS+IFNg or carrier (PBS) (n=6-7). **(B)** *Ass1* expression in microglia sorted from *Acod1*^-/-^ and wt mice treated for 16 h with PBS or LPS (3 mg/kg) (n=6 mice per group). **(C)** *Ass1* and *Il-1b* expression in primary *Acod1*^-/-^ and wt microglia transfected for 24 h with si*Ass1* or siCtrl and then treated with LPS+IFNg for another 24 h (n=8-10). **(D)** mRNA expression of *Il-1b, Sall1, Tmem119, Cx3cr1, Trem2* and *Mertk* in microglia sorted from wt mice treated with for 3 h with argininosuccinate (ASA) (50 mg/kg) and then for 4 h with LPS (1 mg/kg) (n=7-8 mice per group). **(E)** GSEA for polyamine metabolism-related genes based on bulk RNAseq analysis in microglia sorted form LPS-treated *Acod1*^-/-^ and wt mice (n=3 mice per group). **(F)** Spermidine amounts in primary *Acod1*^-/-^ and wt microglia treated for 24 h with PBS or LPS+IFNg (n=9-10). **(G)** *Odc1* and *Il-1b* expression in primary wt microglia transfected for 48 h with 30 nM si*Odc1* or siCtrl and then treated for 4 h with LPS+IFNg (n=9). **(H)** *Il- 1b* expression in primary *Acod1*^-/-^ microglia after treatment for 24 h with PBS, spermidine (SPD), LPS+IFNg or respective controls (n=11). *p < 0.05, **p < 0.01, ***p<0.001, ****p<0.0001. n.d.: non-detectable

In inflammatory macrophages iNOS and ARG1 compete to convert arginine to citrulline and ornithine, respectively (36). While citrulline is the substrate for ASS1, ornithine is metabolized by ornithine decarboxylase 1 (ODC1) to putrescine; the latter is the rate-limiting reaction for the synthesis of polyamines, spermidine and spermine (36). Polyamines suppress inflammation and facilitate resolution of inflammation (36). In accordance with increased argininosuccinate production, gene expression related to polyamine metabolism was downregulated in microglia of *Acod1^-/-^* LPS-treated mice (Figure 3E). Accordingly, spermidine levels were decreased in *Acod1^-/-^* microglia treated with LPS+IFNg (Figure 3F). *Odc1* siRNA silencing (despite its low deletion efficiency) increased *Il-1b* expression in inflammatory microglia (Figure 3G). Moreover, spermidine reduced *Il-1b* expression in inflammatory *Acod1^-/-^* microglia (Figure 3H).

### ACOD1 regulates arginine metabolism via regulation of ACLY

We also observed increased citrate levels in LPS+IFNg-treated microglia, while this effect was repressed in ACOD1 deficient microglia (Figure 4A). Citrate is transported via SLC25A1 from the mitochondria to the cytoplasm, where it is converted by ACLY to acetyl-CoA (45). ACLY-mediated acetyl-CoA production is upregulated in inflammatory macrophages facilitating the induction of inflammatory gene expression by fostering histone acetylation (46). Microglia of LPS-treated *Acod1*^-/-^ mice displayed increased *Acly* expression compared to microglia from wt counterparts (Figure 4B). In accordance, *Acly* and *Slc25a1* expression was upregulated in *Acod1*^-/-^ compared to wt primary microglia treated with LPS+IFNg (Figure 4C). ACLY phosphorylation was enhanced in *Acod1*^-/-^ compared to wt LPS+IFNg-treated microglia 2 h after citrate application (Figure 4D). Consequently, total acetyl-CoA amounts were elevated in *Acod1*^-/-^ compared to wt microglia treated with LPS+IFNg (Figure 4E). Moreover, the ACLY inhibitor BMS303141 abolished *Il-1b* expression in LPS+IFNg-treated microglia (Figure 4F). Interestingly, BMS303141 also decreased *Ass1* expression and increased *Odc1* and spermidine synthase (*Srm*) expression in LPS+IFNg-treated microglia (Figure 4G). These data show that ACOD1 deficiency in microglia promotes ACLY activity, which fosters reprograming of arginine metabolism and promotes the inflammatory response (Figure 4H) (46–48).

**Figure 4.**
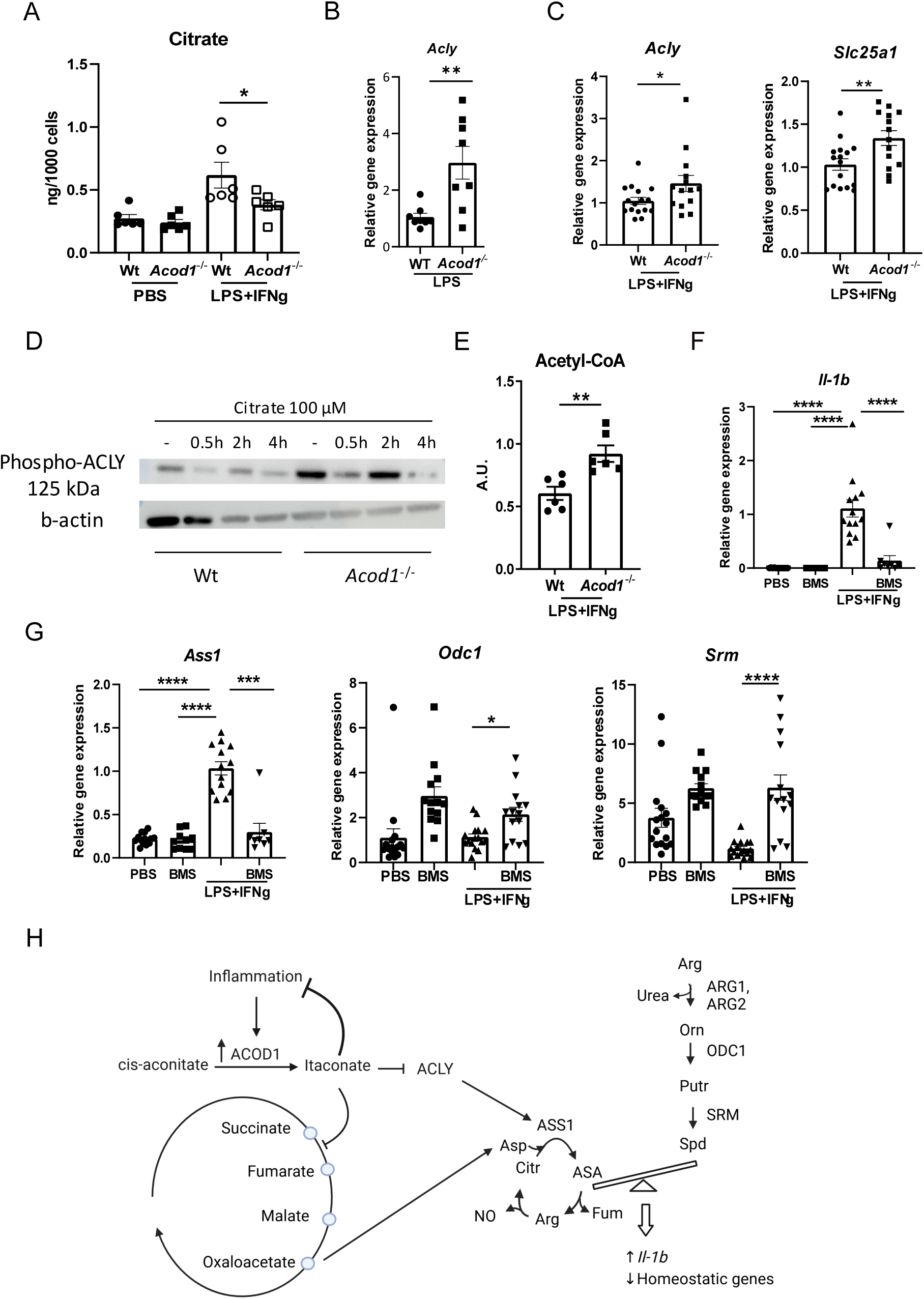
ACOD1 regulates arginine metabolism via ACLY. **(A)** Citrate amounts in *Acod1*^-/-^ and wt microglia treated for 24 h with PBS or LPS+IFNg (n=6). **(B)** *Acly* expression in microglia sorted from *Acod1*^-/-^ and wt mice treated for 16 h with LPS (3 mg/kg) (n=8 mice per group). **(C)** *Acly* and *Slc25a1* mRNA expression in *Acod1*^-/-^ and wt microglia treated for 24 h with LPS+IFNg (n=14-16). **(D)** Western blot analysis for phospho-ACLY after overnight treatment of *Acod1*^-/-^ and wt microglia with LPS+IFNg followed by treatment or not with 100 μM citrate for 0.5 h, 2 h or 4 h using b-actin as a loading control. **(E)** Acetyl-CoA levels in *Acod1*^-/-^ and wt primary microglia treated for 24 h with LPS+IFNg (n=6). **(F)** *Il-1b* mRNA expression in *Acod1^-/-^* microglia treated for 24 h with BMS303141 (20 μM), LPS+IFNg or respective controls (n=8-13). **(G)** *Ass1*, *Odc1* and *Srm* expression in *Acod1^-/-^* microglia after treatment for 24 h with PBS, BMS303141, LPS+IFNg or respective controls (n=8-14). **(H)** ACOD1 regulates via ACLY, arginine metabolism and the inflammatory response in microglia. *p < 0.05, **p < 0.01, ***p<0.001, ****p<0.0001.

## Discussion

The ACOD1-itaconate axis has emerged as a significant regulator of inflammation, as demonstrated by numerous studies in macrophages (2, 5, 6). However, less is known about its role in microglia, the resident macrophage-like cells of the brain. We confirmed that similarly to macrophages, microglia upregulate ACOD1 expression in response to inflammation, which provides negative feedback on the inflammatory response. LPS was identified as the most potent inducer of ACOD1 expression among different TLR ligands and cytokines. Moreover, *Acod1* and itaconate were selectively upregulated by LPS in microglia among glial and neuronal cells in the mouse brain.

Some recent studies have addressed the role of ACOD1 in microglia. Traumatic brain injury (TBI) in mice triggered *Acod1* expression in microglia (49). Microglia-specific ACOD1 deficiency exacerbated TBI-associated inflammation, neurodegeneration and neurological dysfunction and distorted microglial bioenergetics, while 4-OI restored microglial oxidative metabolism (49). In a model of intracerebral hemorrhagic stroke, microglia-specific ACOD1 deficiency reduced erythrocyte clearance thereby aggravating disease, while itaconate or 4-OI restored microglial phagocytic capacity (50). In injury-induced brain ischemia ACOD1 deficient mice displayed aggravated neuroinflammation, blood-brain barrier disruption and brain injury (51). In spinal cord injury, *Acod1* expression was elevated in spinal cords, while overexpression of *Acod1* or treatment with itaconate led to suppression of LPS-induced inflammation in microglia (52). 4-OI and dimethyl itaconate (DMI) were suggested to halt experimental autoimmune encephalomyelitis (EAE) progression in mice (53, 54), although the ameliorating effect of 4-OI in EAE was disputed by others (55). Moreover, *Acod1^-/-^* deficient mice presented greater microglia density and allograft inflammatory factor 1 (AIF1) reactivity in the CA1 region and dentate gyrus upon systemic LPS stimulation (56). Accordingly, systemic DMI treatment restrained microgliosis in the hippocampus of *Toxoplasma gondii*-infected mice (57).

Here, we demonstrate the anti-inflammatory role of the ACOD1-itaconate axis by RNA-seq in microglia sorted from brains of *Acod1^-/-^* LPS-treated mice. *Acod1^-/-^* microglia showed enhanced inflammation, manifested by increased IL-6 and IL-1b production. Furthermore, we explored the immunometabolic role of ACOD1 in microglia. ACOD1 deficiency enhanced ACLY expression and acetyl-CoA amounts. Increased ACLY activity in LPS-treated macrophages was previously shown to promote *Il-6* and *Il-1b* expression via upregulation of histone acetylation in their promoter regions (46, 59). In accordance, itaconate-bearing lipid nanoparticles targeting atherosclerotic plaques reduced H3K27ac marks in inflammatory genes in myeloid cells (60). We show that increased ACLY activity in *Acod1^-/-^* microglia upregulates *Ass1* expression and argininosuccinate amounts. Furthermore, itaconate depletion in *Acod1^-/-^* microglia lifts SDH inhibition, leading to the unimpeded metabolic pathway fumarate – malate – aspartate – argininosuccinate, fueling the aspartate-argininosuccinate shunt (42). Hence, increased substrate availability combined with upregulated ASS1 expression in *Acod1^-/-^* microglia may foster the aspartate-argininosuccinate shunt, thereby sustaining arginine regeneration and nitric oxide production (36, 37, 42). The ASS1-argininosucciante axis promoted *Il-1b* expression, standing in accordance with previous studies showing that ASS1-mediated depletion of citrulline, which inhibits JAK2-STAT1 signaling, is required for host defense against bacterial infection (41). Accordingly, previous reports showed that iNOS inhibits inflammasome activation and promotes inflammasome tolerance in synergy with itaconate in LPS-treated macrophages (61). Finally, our data suggest that tilting arginine metabolism towards argininosuccinate production downregulates biosynthesis of polyamines. Polyamines mediate anti-inflammatory and pro-resolving effects (62–65); hence, impediment of polyamine synthesis favors the proinflammatory phenotype of ACOD1 deficient microglia. Taken together, these findings demonstrate that the ACOD1/itaconate axis maintains balance between argininosuccinate and polyamine synthesis, thereby revealing a novel immunometabolic mechanism regulating microglial inflammatory responses.

## Acknowledgments

We thank Christine Mund, Denise Kaden and Catleen Conrad for technical assistance.

## Funding

This work was supported by grants from the Deutsche Forschungsgemeinschaft (AL 1686/6-1, SFB-TRR 205 project A07, BR 4853/2-1 and IRTG3019 project P02 to VIA, and a major instrument grant support (INST 269/910-1 FUGG) to MP).

## Data availability

RNA-Seq data are available in: https://www.ncbi.nlm.nih.gov/geo/query/acc.cgi?acc=GSE299665.

## Conflict of Interest

The authors have no conflicts of interest.

## Author contributions

EK: Investigation, Data Curation, Visualization, Writing - Original Draft; CY: Investigation, Data Curation, Visualization; AS: Data Curation; GF: Investigation; SD: Investigation; NN: Investigation; ST: Methodology, Investigation, Data Curation, NZ: Methodology, Data Curation, Resources; BW: Project administration; PV: Resources, Project administration MP: Methodology, Investigation, Resources; TC: Conceptualization, Resources, Writing - Review & Editing; VIA: Conceptualization, Resources, Writing - Review & Editing; Supervision, Project administration, Funding acquisition.

## Notes

### Competing Interest Statement

The authors have declared no competing interest.

### Summary of Updates

Funding source added: major instrument grant support (INST 269/910-1 FUGG) to MP Data availability RNA-Seq data are available in: https://www.ncbi.nlm.nih.gov/geo/query/acc.cgi?acc=GSE299665.

