## Supplemental data for "ACOD1 regulates microglial arginine metabolism and inflammatory responses"

**Supplementary data**

**Supplementary Figure 1. LPS induces *Acod1* expression in primary microglia. (A-C)** RNA-seq in primary microglia treated for 4 h with LPS or carrier (PBS) (n=4). **(A)** Volcano plot showing differentially expressed genes. **(B)** GSEA for inflammatory response-related genes. **(C)** Heatmap of differentially expressed TCA cycle-related genes (padj < 0.05). **(D)** *Acod1* mRNA expression in primary microglia cells treated for 4 h with LPS+IFNg or carrier (PBS) (n=6). **(E)** Western blot analysis for ACOD1 after treatment of primary microglia cells for 24 h with PBS or LPS+IFNg, using Vinculin as a loading control. **p < 0.01.

**Supplementary Figure 2. Expression of *Acod1* in different cell types. (A)** *Acod1* mRNA expression in $CD45^{high}CD11b^{+}Ly6G^{-}$ cells and neutrophils sorted from bone marrow of wt mice and treated in vitro for 4 h with LPS (100 ng/ml) or carrier (PBS) (n=4 per group). **(B)** *Acod1* mRNA expression in sorted $CD4^{+}$ cells, $CD8^{+}$ cells and $CD45^{high}CD11b^{+}Ly6G^{-}$ cells isolated from spleens of mice treated for 16 h with LPS (3 mg/kg) or PBS (n=5 per group). **(C)** Itaconate amounts in microglia and $CD45^{high}CD11b^{+}Ly6G^{-}$ cells sorted from spleens of mice treated for 16 h with LPS (3 mg/kg)(n=2). **(D)** Number of sorted microglia and monocytes/macrophages from brains of mice treated for 16 h with LPS (3 mg/kg). **(E)** Schematic depiction of sequential isolation of microglia, oligodendrocytes, astrocytes and neurons from mouse brains. **(F)** *Acod1* expression in microglia sorted from brains of wt and *Acod1*$^{-/-}$ mice treated for 16 h with LPS (3 mg/kg) or PBS (n=6 mice per group). **(G)** Itaconate amounts in primary wt and *Acod1*$^{-/-}$ microglia treated for 24 h with LPS+IFNg or carrier (PBS) (n=6 per group). *p < 0.05,**p < 0.01.

**Supplementary Figure 3. Effect of different TLR ligands and cytokines on *Acod1* expression in microglia.** *Acod1* mRNA expression in BV2 cells treated or not for 4 h with various TLR ligands (n=6) **(A)** or cytokines (n=7) **(B)**. *p < 0.05, **p < 0.01, ***p < 0.001, ****p < 0.0001.

**Supplementary Figure 4.** Succinate and succinate/fumarate in primary wt and *Acod1*$^{-/-}$ microglia treated for 24 h with PBS or LPS+IFNg (n=6), **p < 0.01.

**Table 1. Primer sequences**

| Gene name | Primer | Sequence (5' to 3') |
|---|---|---|
| *18s* | forward | GTTCCGACCATAAACGATGCC |

| | | |
|---|---|---|
| *18s* | reverse | TGGTGGTGCCCTTCCGTCAAT |
| *Acod1* | forward | CTCCCACCGACATATGCTGC |
| *Acod1* | reverse | GCTTCCG TAGAGCTGTGA |
| *Il-1b* | forward | TGGGATGATGATGATAACCTGC |
| *Il-1b* | reverse | TCGTTGCTTGGTTCTCCTTGTA |
| *Il-6* | forward | CCTTCCTACCCCAATTTCCAAT |
| *Il-6* | reverse | AACGCACTAGGTTTGCCGAGTA |
| *Ass1* | forward | CTGCTATTCACTGGCACCCC |
| *Ass1* | reverse | GATCATTTCGGCCCTTGAACC |
| *Sall1* | forward | GCTTGCACTATCTGTGGAAGAGC |
| *Sall1* | reverse | CTGGGAACTTGACAGGATTGCC |
| *Tmem119* | forward | GTGTCTAACAGGCCCCAGAA |
| *Tmem119* | reverse | AGCCACGTGGTATCAAGGAG |
| *Cx3cr1* | forward | AGGACACAGCCAGACAAG |
| *Cx3cr1* | reverse | TCAGGGGAGAAAGCAAG |
| *Trem2* | forward | GTACTGGTGGAGGTGCTGGA |
| *Trem2* | reverse | GGAGGTGCTGTGTTCCACTT |
| *Mertk* | forward | CGGTAATAATCACCACTGTAAATCTTTCT |
| *Mertk* | reverse | TTGCGGGATGACATGACTGT |
| *Odc1* | forward | CGCAGTCAAGTGTAACGATAGC |
| *Odc1* | reverse | GAGACTTGTTTACAAGGATTTGCAT |
| *Acly* | forward | AGGAAGTGCCACCTCCAACAGT |
| *Acly* | reverse | CGCTCATCACAGATGCTGGTCA |
| *Slc25a1* | forward | GGAGGCACACAAATACCGGA |
| *Slc25a1* | reverse | GGTGCCCTTGTAGAATGCCT |
| *Srm* | forward | AGGAGATGATCGCCAACCT |
| *Srm* | reverse | TTCACCACTTCCCGTAGGAC |

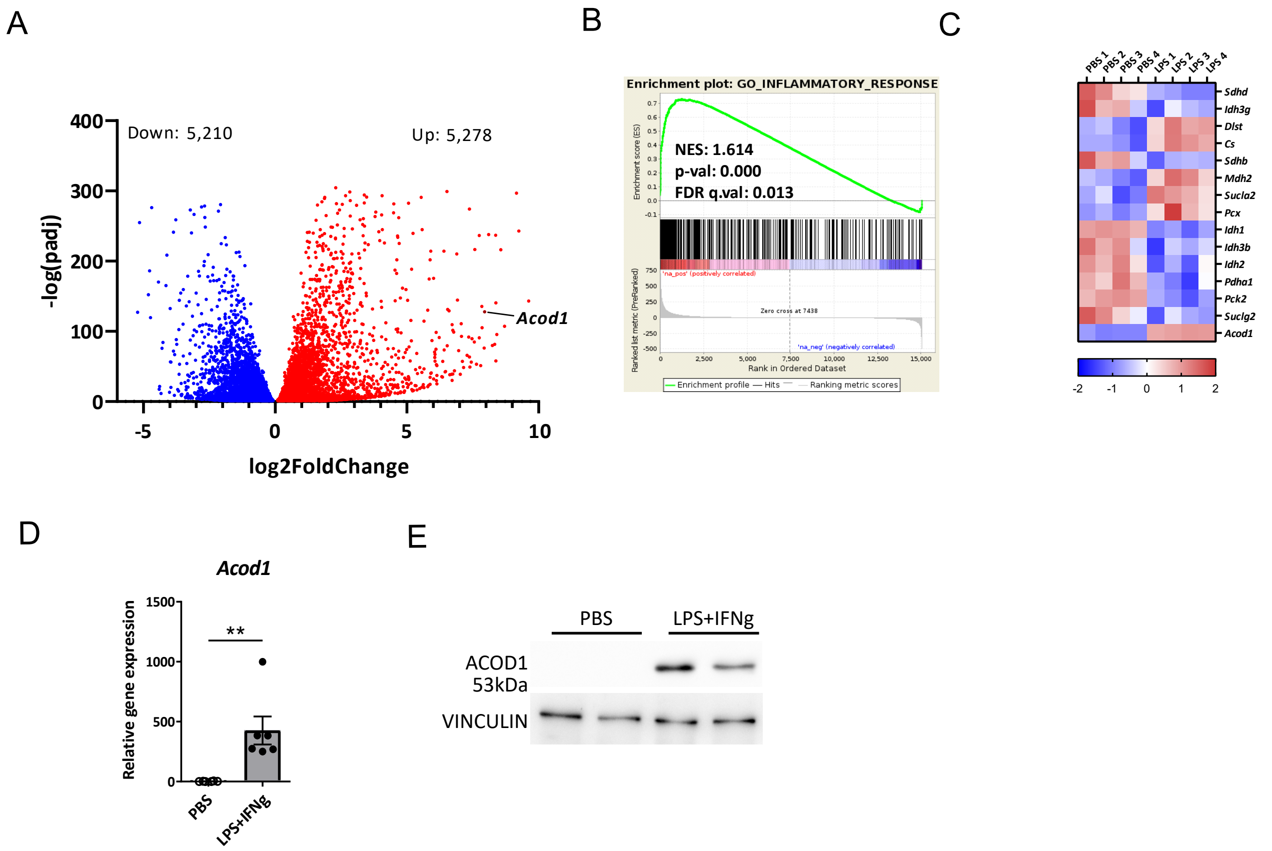
A
Down: 5,210
Up: 5,278
-log(padj)
log2FoldChange
Acod1
B
Enrichment plot: GO_INFLAMMATORY_RESPONSE
NES: 1.614
p-val: 0.000
FDR q.val: 0.013
Enrichment score (ES)
Ranked list metric (PreRanked)
Zero cross at 7438
Rank in Ordered Dataset
Enrichment profile
Hits
Ranking metric scores
C
PBS 1
PBS 2
PBS 3
PBS 4
LPS 1
LPS 2
LPS 3
LPS 4
Sdhd
Idh3g
Dlst
Cs
Sdhb
Mdh2
Sucla2
Pcx
Idh1
Idh3b
Idh2
Pdha1
Pck2
Suclg2
Acod1
D
Acod1
Relative gene expression
**
PBS
LPS+IFNg
E
PBS
LPS+IFNg
ACOD1
53kDa
VINCULIN

Figure S1

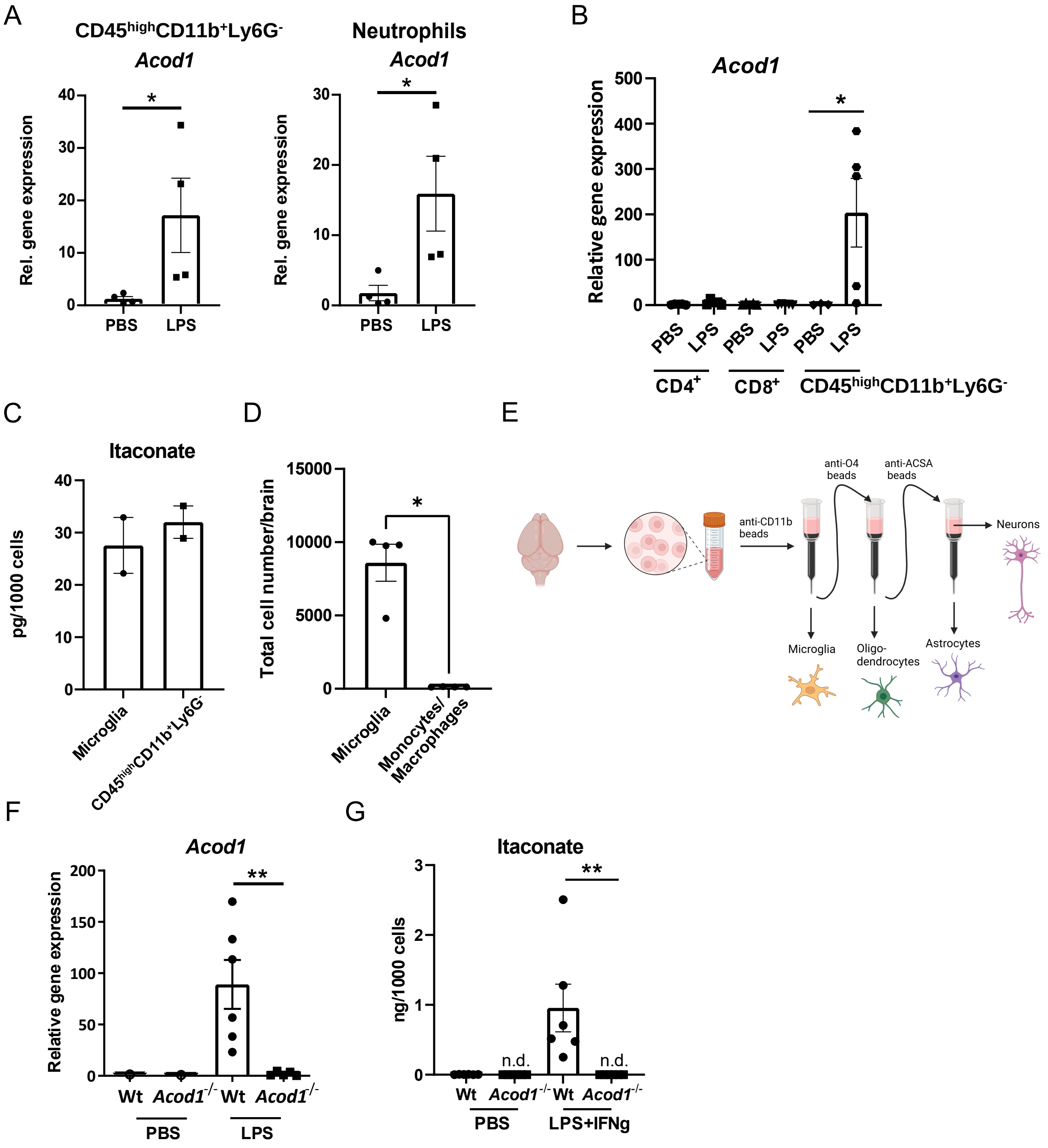
A
CD45highCD11b+Ly6G-
Acod1
Rel. gene expression
PBS
LPS
Neutrophils
Acod1
B
Acod1
Relative gene expression
CD4+
CD8+
CD45highCD11b+Ly6G-
C
Itaconate
pg/1000 cells
Microglia
CD45highCD11b+Ly6G-
D
Total cell number/brain
Microglia
Monocytes/ Macrophages
E
anti-CD11b beads
anti-O4 beads
anti-ACSA beads
Neurons
Microglia
Oligo-dendrocytes
Astrocytes
F
Acod1
Relative gene expression
Wt
Acod1-/-
PBS
LPS
G
Itaconate
ng/1000 cells
n.d.
LPS+IFNg

Figure S2

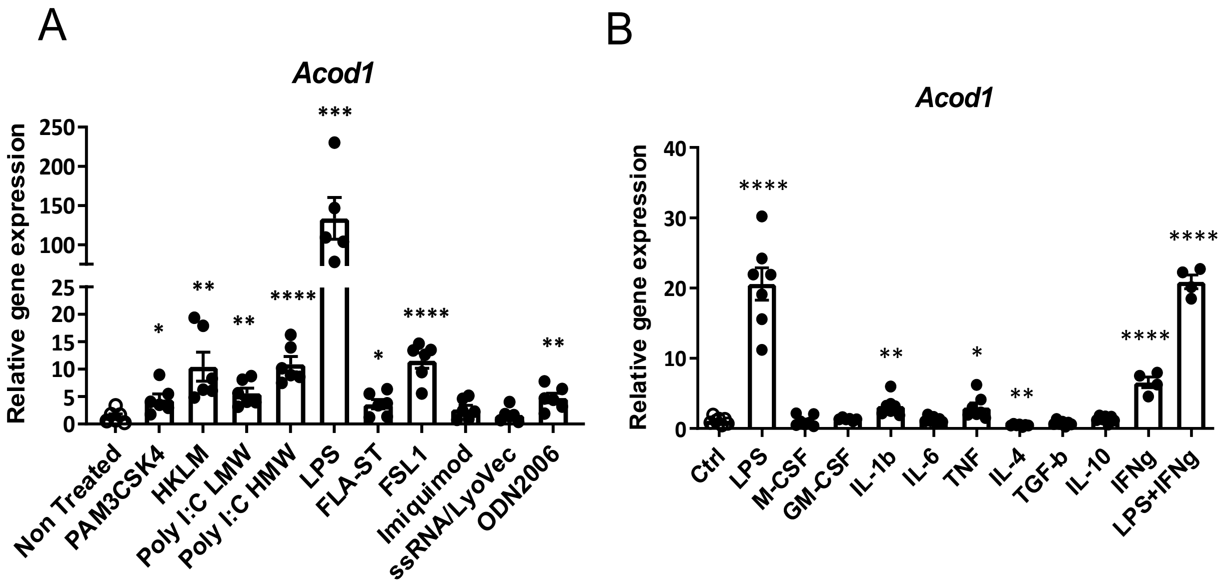
A
Acod1
Relative gene expression
250
200
150
100
25
20
15
10
5
0
*
**
**
****
***
*
****
**
Non Treated
PAM3CSK4
HKLM
Poly I:C LMW
Poly I:C HMW
LPS
FLA-ST
FSL1
Imiquimod
ssRNA/LyoVec
ODN2006
B
Acod1
Relative gene expression
40
30
20
10
0
****
**
*
**
****
****
Ctrl
LPS
M-CSF
GM-CSF
IL-1b
IL-6
TNF
IL-4
TGF-b
IL-10
IFNg
LPS+IFNg

Figure S3

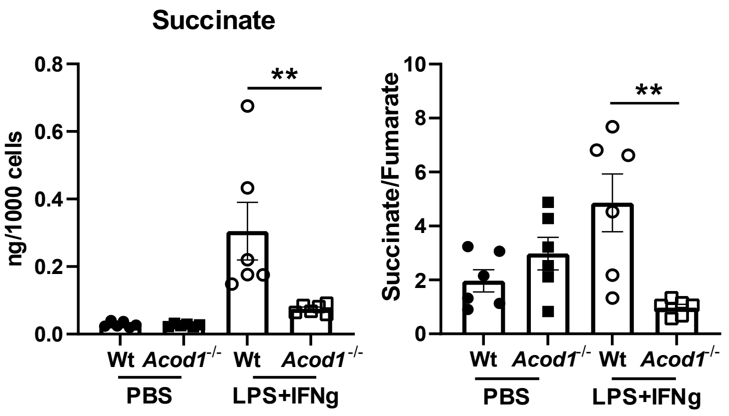
Succinate
ng/1000 cells
0.8
0.6
0.4
0.2
0.0
**
Wt
Acod1-/-
Wt
Acod1-/-
PBS
LPS+IFNg
Succinate/Fumarate
10
8
6
4
2
0
**
Wt
Acod1-/-
Wt
Acod1-/-
PBS
LPS+IFNg

Figure S4
